# Stochastic modelling of prostate progenitor architecture

**DOI:** 10.1101/2025.06.19.660586

**Authors:** Christo Morison, Esther Baena, Weini Huang

**Author notes:** joint supervision (Esther Baena, Weini Huang). corresponding authors (C.M.), (W.H.).

## Abstract

Prostate cancer is a significant global health concern. Experimental and clinical research have contributed to the improvement of treatment outcomes dramatically in the past decade. However, much of the structure of the prostate and its constituent cell types remains unclear: observations of phenotypic switching between basal and luminal cells, as well as intermediate cell types between them, make classifying the components of the prostate an ongoing challenge. Importantly, a tumour’s cell type of origin seems to play a role in the aggressiveness and genetic heterogeneity of the cancer. To understand how different structures between cell types impact the emergence and treatment of prostate cancer, we model the emergence of a tumour by a fit mutant arising amongst healthy cells via a compartmental modelling approach. We label cells in different compartments based on their phenotype and allow for phenotypic switching between them. We measure the time until and probability of invasion and extinction of this mutant in healthy prostates through individual-based simulations. We find that less-tight population size regulation of the luminal compartment allows for the explosive population growth associated with cancer. Tumours arising in the basal compartment require more time to spread but are more strongly embedded within the prostate, explaining the longer latency and higher aggressiveness and persistence associated with these tumours. Interestingly, the inclusion of a hybrid compartment does not qualitatively change the observations, raising questions as to what mechanistic role intermediate phenotypes play in prostate cancer emergence, if any. Our model contributes to dissecting the relationship between prostate structure and outcomes for cancers arising therein.

## Introduction

Prostate cancer is the fourth most commonly diagnosed cancer worldwide and second only to lung cancer among males, accounting for 7.3% of new cancer incidences (over 1.4 million) in 2022 [1]. Affecting predominantly men advanced in age, approximately 80% of cases are diagnosed with non-metastatic, organ-contained disease [1]. While the advent of prostate-specific antigen as a disease marker resulted in a surge of cases in the 1990s and harmful overdiagnosis and overtreatment [2–4], patients with localised disease are diagnosed and treated early with good outcomes including an overall survival rate of 99% for 10 years [5].

The prostate’s primary function is to produce seminal fluid that maintains sperm viability along the ejaculatory duct and urethra [6]. Anatomically, the organ has five regions: the central, transition and peripheral zones, along with the periurethral gland and fibromuscular regions [5]. The tissue in each region is composed of basal and luminal cells, whose population ratio in humans is roughly equal, as well as a small population of neuroendocrine cells [7]. These are distinguishable histologically, as well as via their positioning: basal and neuroendocrine cells encircle luminal cells, which enclose the lumen and secretions therein [5]. Figure 1 depicts a simplified sketch of the prostate.

**Figure 1.**
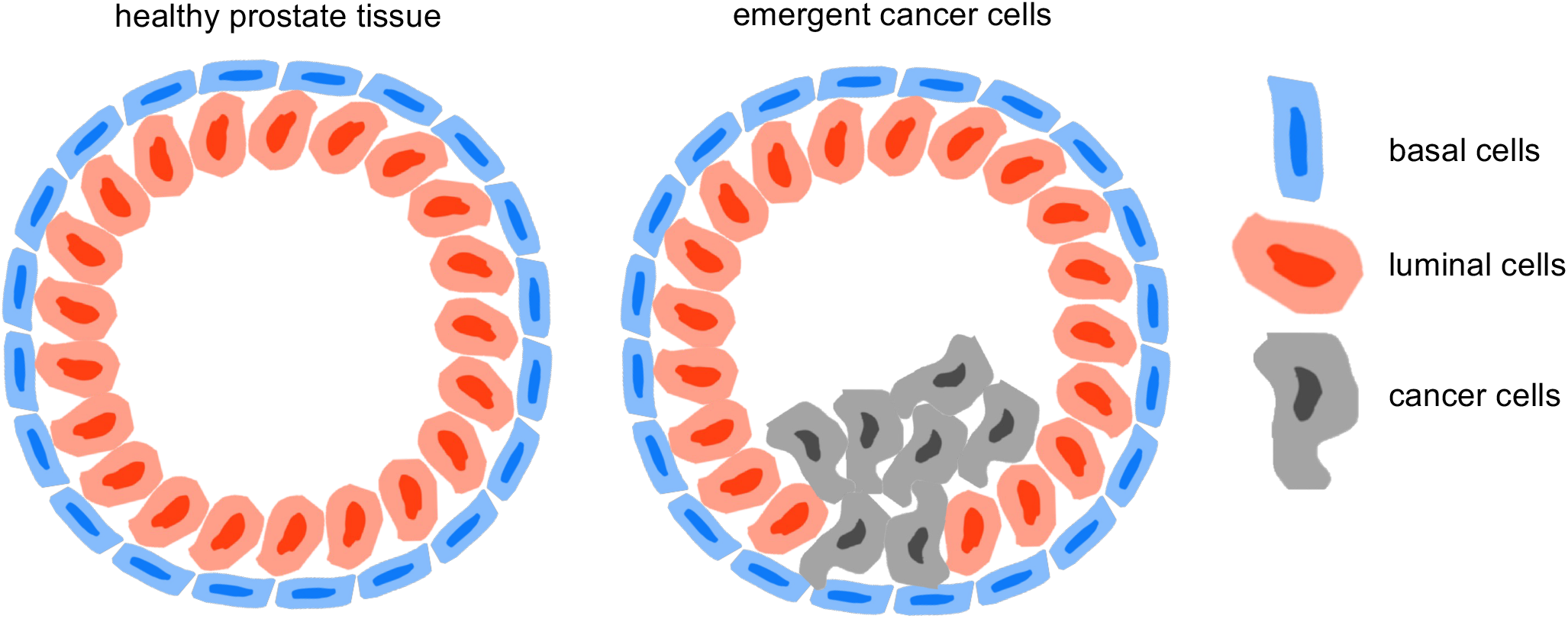
Sketch of the two primary cell types in the prostate: basal (blue) cells ring luminal (red) cells, enclosing the prostate duct. The right panel depicts the arrival of cancer (grey) cells, which may spill into the duct before possibly exiting the prostate altogether. Figure adapted from Rebello et al. [5].

Tumours often occur in the peripheral region, most distant from the urethra [5]. However, while the cell that originates the cancer can be of any type, when detected, cancers are predominately luminal, rather than basal [8]. Importantly, it has been observed that tumours derived from luminal cells have shorter tumour latency, the time from the cancer’s initiation to its detection [9, 10]. In addition, basal-derived tumours are of a higher grade and more likely to be aggressive and spread [11, 12]. This suggests that the delays undergone, which result in the higher latency, may be connected to subsequent increased invasiveness. The gene signature associated with these particularly aggressive cancers has also been linked to multipotency in basal cells [13, 14]. As a consequence, understanding the structure between cell types in the prostate may illuminate the role of the cell of origin in determining the aggressiveness of prostate tumours.

Both basal and luminal types have progenitor capabilities and can produce mature cells via differentiated cell division as well as self-proliferate to generate new progenitor cells [15]. However, there is no consensus as to how many basal and luminal (sub)types exist [16]. Additionally, how these cell types are related to one another—that is, how they may transition or differentiate between states—is unclear. While phenotypic switching between cell types has been observed in situations such as wound healing, where e.g. depletion in the luminal compartment is replenished by basal progenitors [16], evidence of hybrid progenitor types (with both basal and luminal characteristics) is increasing [15, 17, 18].

Compartmental models are commonplace in mathematical biology, with uses ranging from labelling members of population during an epidemic [19] to labelling cell phenotypes during subsequent rounds of differentiation in haematopoiesis [20]. To produce mature blood cells, the progeny of haematopoietic stem cells undergo a differentiation process called lineage commitment, wherein the potential to produce all cell types is progressively lost [20]. In epidemiological models, individuals falling under different labels, e.g. susceptible, infected and recovered, are often stratified into compartments to study population dynamics and disease spread [19]. The compartments ascribed to these populations may be somewhat artificial; nevertheless, by doing so we can use models to study how the architecture of the system affects its dynamics.

Here, we model the prostate by focussing on four cell types: basal, hybrid and two luminal subtypes. The number of luminal subtypes was chosen for two primary reasons: first, because most recent classifications concur on there being at least two subtypes [16]; second, to keep the system fairly simple, we abstained from choosing a large or variable number. With the analysis for only two luminal subtypes, we will be able to draw some conclusions about what happens if more exist, since the labels are less important than the distinctions between the compartments (such as how they are connected to one another, or what their population sizes are, etc.). On the other hand, the singular basal type is in agreement with many (murine) studies [16, 17, 21–24].

A subset of basal and luminal cells maintains a homeostatic population of mature cells through differentiation, and four types (basal, hybrid and two luminal subtypes) are connected via phenotypic switching. We use stochastic simulations to study the impact of different structures between these cell types on the emergence and treatment of prostate cancer.

### Model

We consider a compartmental model of the prostate with two types of compartments: progenitor compartments, which can produce cells of their own type (self-renewal) and mature compartments (which cannot). As depicted in Figure 2, there are four progenitor compartments, named HP for hybrid (purple), BP for basal (blue), LP_1_ for luminal 1 (red) and LP_2_ for luminal 2 (pink). Self-renewing is depicted by green arrows; phenotypic switching between different types of progenitor cells (compartments) is depicted by yellow arrows. The three basal and luminal progenitor compartments (BP, LP_1_ and LP_2_) have corresponding mature compartments, denoted by the same colour with black centres (B, L_1_ and L_2_, respectively), where differentiation is depicted by black arrows. We denote the population sizes of the compartments at time *t* by the name of the compartment itself: for instance BP(0) is the initial population size in the basal progenitor compartment, and L_1_(10) is the population size in the luminal 1 mature compartment at time *t* = 10.

**Figure 2.**
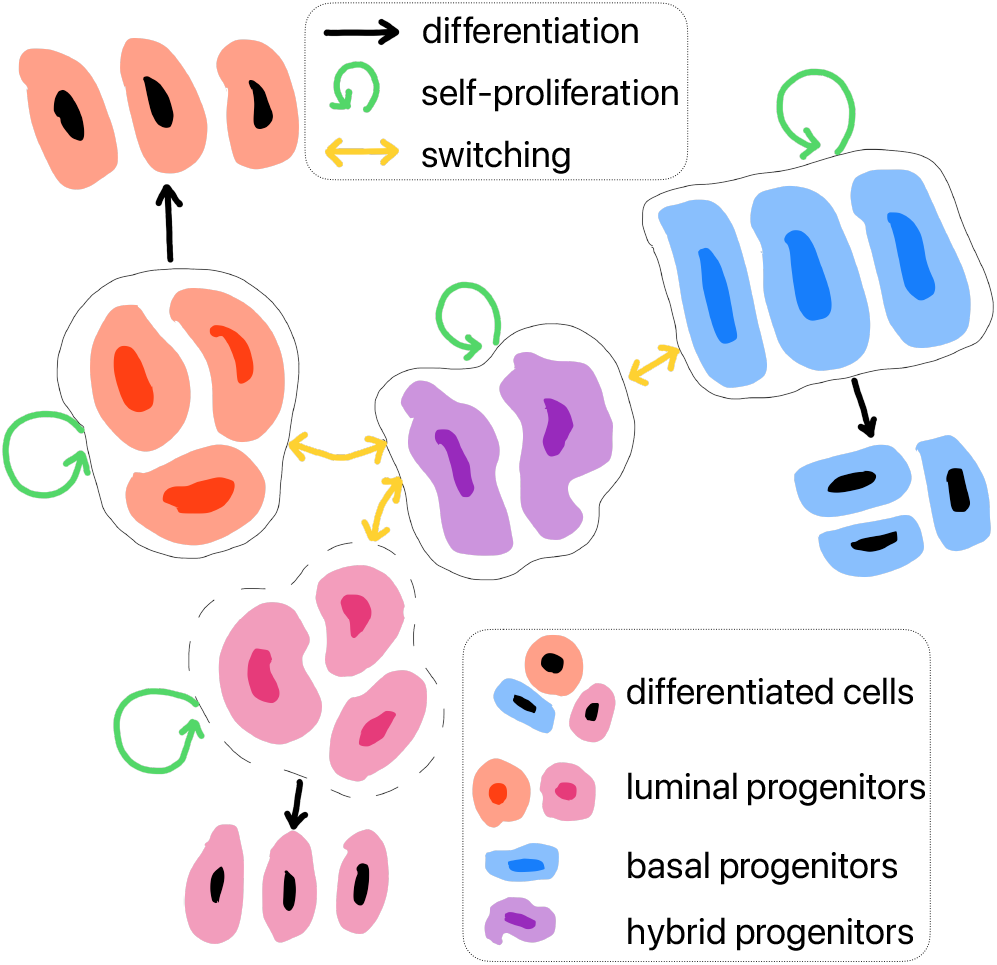
Compartmental model of prostate architecture: hybrid/HP (purple), basal/BP (blue), luminal 1/LP_1_ (red) and luminal 2/LP_2_ (pink) cells can self-proliferate (green arrows), differentiate (black arrows) or switch phenotypes (yellow arrows). Progenitor compartments have tight population size regulation (solid lines) or not (dashed lines); in particular, in our model, LP_2_ has a looser population size regulation than the other compartments.

### Compartment dynamics

Each progenitor compartment undergoes a Moran Birth-death process at some prescribed initial population size [25]. First, a cell is selected, with a probability depending on its fitness, to divide. Divisions can be symmetric, producing two identical daughter cells of the same type (wild-type, representing normal cells, and fit mutants, representing tumour cells). Or, they can be asymmetric, where the division results in either differentiation or phenotypic switching with different rates: one daughter cell resembles the mother cell, and the other is mature or of a different progenitor type, respectively. A cell is then killed randomly (without regard for cell fitness) in the compartment that has increased in population, leaving the population size constant. This is to mimic regulated tissues before progressing to aggressive tumours.

### Invasion and extinction

Compartments are initially assumed to be filled with wild-type (normal) cells with unit fitness, save for one mutant cell with fitness 1 + *s* in the “seeded” compartment, representing a malignant tumour cell. Note that here the rate of a birth event (which, as described in the previous section, also induces a death event) is proportional to the fitness of the individual. The mutant cell and its descendants all have self-proliferative capabilities and can switch phenotypes in a manner identical to wild-type cells. The mutant is considered to have invaded when the mutant population of luminal type reaches *r*_inv_ = 40% of the initial, homeostatic, wild-type luminal population (sum of LP_1_, LP_2_, L_1_ and L_2_). This threshold was chosen as an approximation of the size of tumours upon being detectable; however, since the parameters used in this study are in arbitrary units, this is not critical, and another nearby value could have been chosen with similar results. The time for this invasion to occur (conditioned on invasion occurring), which can also be thought to be the tumour latency, is written *T*_inv_, measured here in arbitrary units. The probability of invasion occurring is *p*_inv_, the value of which is less than one, despite the mutants having a greater fitness than wild-type cells. This is due to stochastic effects, which are appreciable for small mutant population sizes and lead to mutant extinction for some realisations. The probability that the mutant goes extinct before invading is written *p*_ext_, with time of extinction (conditioned on extinction occurring) *T*_ext_; there is also a possibility that the mutant does not invade nor go extinct before the stopping time *T*_end_ of the simulation. By simulating many stochastic realisations of the model, we can estimate *p*_inv_ and *p*_ext_ for specific parameter sets, along with expected values of *T*_inv_ and *T*_ext_. All simulations were implemented via a standard Gillespie algorithm [26].

### Model comparison

In line with experimental and clinical observations, our model has three distinct features. First, looser population size regulation in one of the luminal compartments, in correspondence with most cancers being of luminal, rather than basal, type [8]. In our model, LP_2_ differs from the other progenitor compartments in its population size regulation: to model the possibility of looser regulation, we allow for mutant cells to avoid the death event with probability *N*_m_/(*N*_wt_ + 2*N*_m_), for *N*_wt_ and *N*_m_ the LP_2_ wild-type and mutant population sizes, respectively. This probability increases from 0, when there are no mutant cells, to 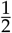, when mutants are fixed in the compartment. This feature of the model will be referred to as looser regulation of LP_2_, written *L* = 1. When *L* = 0, LP_2_ is treated as the other progenitor compartments, with strict Moran Birth-death dynamics.

Second, the presence of a hybrid progenitor compartment, HP. As previously described, our model consists of four progenitor compartments, labelled HP, BP, LP_1_ and LP_2_, with the latter three having corresponding differentiated compartments (B, L_1_ and L_2_, respectively), and each with varying initial population sizes (see Table 1) [16]. The presence of HP will be written *H* = 1. When *H* = 0, there is no hybrid progenitor compartment; instead, the basal compartment BP connects the two luminal compartments LP_1_ and LP_2_ with phenotypic switching: LP_1_ ↔ BP ↔ LP_2_ is the resulting topology, where ↔ denotes bilateral switching.

**Table 1.**
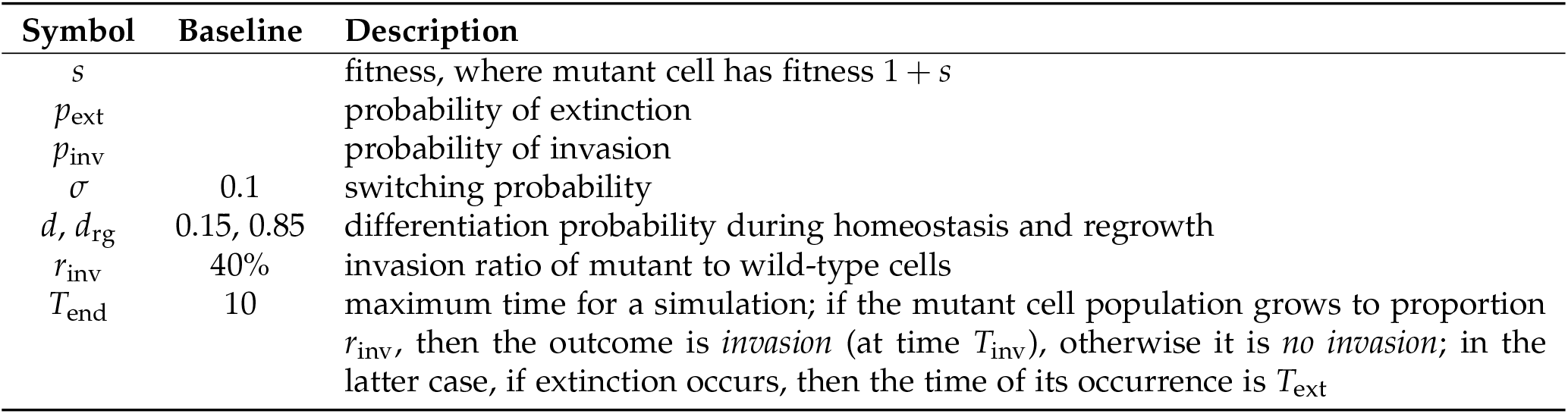
Notation and baseline parameter values, which are used for all simulations unless otherwise specified. Initial population sizes for {HP, BP, LP_1_, LP_2_, B, L_1_, L_2_} are {200, 1000, 100, 900, 10000, 1000, 9000}.

Third, we include phenotypic switching between progenitor types: although most tumours are phenotypically luminal, they can originate in the basal compartment [11, 12]. There is also evidence of the basal compartment helping replenish the luminal compartments when they are depleted due to injury [15]. Inclusion of phenotypic switching will be denoted *S* = 1. When *S* = 0, no switching is permitted between progenitor types; instead, during a progenitor compartment birth event, the resulting daughter cell either stays within the same compartment (self-renewal), or moves into the corresponding mature compartment (differentiation).

We can explore different model structures by including or excluding looser population size regulation of LP_2_ (*L* = 1), the presence of HP (*H* = 1) and the inclusion of phenotypic switching (*S* = 1). We will write this trio of attributes as a triple {LHS} whose entries are 0 or 1. For instance, {111} represents the full model, whereas {101} represents a model with a less-regulated luminal 2 compartment and phenotypic switching, but no hybrid compartment. By comparing {000}, {100}, {101} and {111}, we can measure the impact and importance of each attribute, since they are added one at a time. Note that the full {111} model is used throughout unless otherwise specified.

### Treatment

If the mutant population invades, then we suppose that it is large enough to clinically detect and thus to treat. To represent androgen deprivation therapy (ADT), a common treatment option for both localised and advanced prostate cancers, we assume a depletion of the luminal (LP_1_, LP_2_, L_1_ and L_2_) and hybrid (HP) cell populations [27, 28]. Treatment is modelled to remove a random proportion *r* (between 90% and 95%) of the luminal and hybrid cells [17]; this removal is instantaneous. The realisation continues, with the progenitor cell types undergoing more rapid differentiation (controlled by a regrowth differentiation parameter *d*_rg_ > *d*) to replenish the lost cells.

## Results

The emergence of cancer is modelled by the propensity for a mutant cell to invade the prostate. At low populations, such as those considered here, random events play a significant role in these dynamics: by simulating many realisations of the system, we can characterise the underlying stochastic processes. We first consider a single mutant with fitness 1 + *s* seeded into one of the progenitor compartments (HP, BP, LP_1_ or LP_2_) and inspect the resulting invasion probability and the average invasion time, conditioned on invasion, for the {111} model.

### Invasion versus fitness

Figure 3 demonstrates that invasions are more likely in LP_2_ (which is less tightly regulated) than the other compartments (HP, BP and LP_1_), for which the invasion probability varies similarly with *s*. As the fitness *s* increases, more invasion events take place no matter where the mutant is seeded. The invasion time distinguishes between different compartments, since mutants arising in LP_2_ invade more quickly than those in HP (intermediate invasion times) and LP_1_ and BP (long invasion times). This is due to the fact that switching events must take place in the latter three scenarios before the population can grow in the less-tightly regulated LP_2_: for the HP-seeded mutant, one such switch must occur, whereas two must occur for LP_1_- or BP-seeded mutants. This pattern is exacerbated by an increasing fitness, which itself decreases the invasion time in all cases, since the mutants have a greater advantage in proliferation compared to the wild-type cells. Note that for lower fitnesses, fewer realisations result in invasion, so the invasion times are averaged over fewer instances, hence the leftmost parts of the curves in the right panel of Figure 3 are less smooth; similarly, there are no curves at all for low values of *s* (in particular for HP-, BP- and LP_1_-seeded simulations) in the right panel because no realisations resulted in invasions and thus the invasion time is not defined.

**Figure 3.**
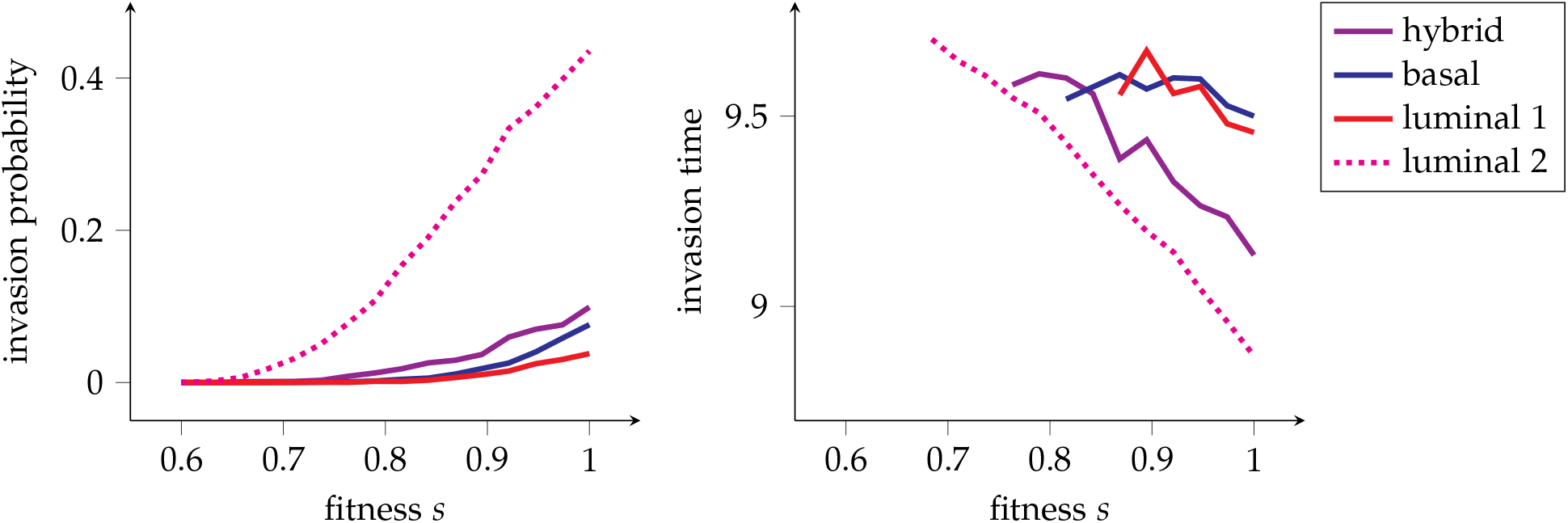
Invasion probability (left panel) and invasion time (right panel; in arbitrary units) versus fitness, averaged over 5500 realisations. Colours denote which compartment was seeded with a single mutant: hybrid/HP (purple), basal/BP (blue), luminal 1/LP_1_ (red) or luminal 2/LP_2_ (pink dotted).

### Invasion versus switching and differentiation

Figure 4 demonstrates the impact of the differentiation and switching probabilities on the invasion probability and time for HP-, BP- and LP_2_-seeded mutants. (See Figure S3 for mutants seeded into LP_1_.) First, note that if the switching probability is zero, then invasion is impossible when HP or BP are seeded. Once above this minimal threshold (which is slightly higher for the BP case, since two switchings must occur compared to one in the HP case; as shown in Figure S3, the LP_1_ case closely follows the BP case), the switching probability plays a limited role; instead, an increase in the differentiation probability both increases the invasion probability and reduces the invasion time. This can be explained with the following reasoning: since mutant cells retain progenitor capabilities (that is, they do not differentiate), they are relatively advantaged when wild-type cells differentiate more. This is because differentiation events do not replace existing mutant cells; therefore, all mutant divisions may replace wild-type cells, whereas only divisions of wild-type cells that result in self-proliferation (rather than in differentiation) can replace mutant cells. Interestingly, the differentiation and switching probabilities influence the invasion time in concert, no matter the seeded compartment, as shown in the right column of Figure 4: the most rapid invasions occur with low switching (so more mutant cell division events result in self-proliferation) and high differentiation (for the aforementioned relative advantage reason) probabilities. This pattern is most evident in the LP_2_ case (which also has the highest invasion probabilities and lowest invasion times), since in this compartment more invasions occur, so the variance is lower.

**Figure 4.**
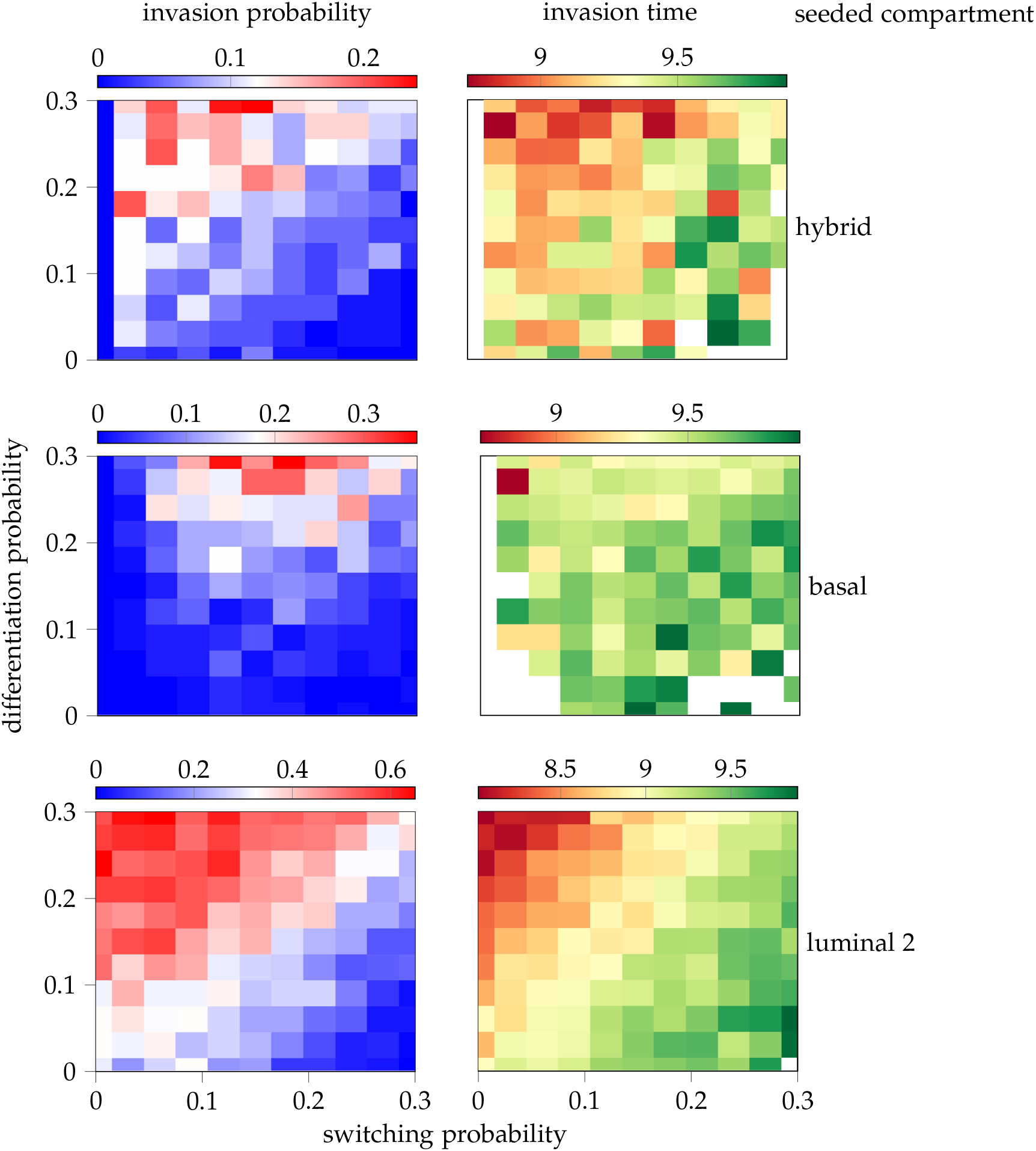
Invasion probabilities (left) and invasion times (right; in arbitrary units) for different switching and differentiation probabilities when the mutant is seeded in the hybrid/HP, basal/BP and luminal 2/LP_2_ compartments. Each cell is averaged over 100 realisations; blank cells in the right column indicate that zero invasions took place.

### Switching and regulation allow for invasion

The invasion probability and the invasion time can also be used to distinguish between our different model systems. For a fixed fitness *s* = 1 (chosen because the patterns of Figure 3 were fairly monotonic in *s*, so a higher fitness would demonstrate underlying behaviours more clearly without corruption), Figure 5 depicts these two quantities for each of the four model types: {000}, {100}, {101} and {111}. First, when LP_2_ is as tightly regulated as the other progenitor compartments, invasion does not occur, as evidenced by invasion probabilities of zero for the {000} model no matter the mutant cell seeding location. If LP_2_ is more loosely regulated, then invasion may occur, though only if the mutant is seeded in that same compartment when no switching is allowed, as in the {100} model. Next, when switching is included, invasion is possible no matter the seeding location (as shown by the {101} model results), though the invasion probability is smaller and the time is longer when the seeding is distant from LP_2_. Finally, when HP is introduced, the model provides similar results to the {101} model, with HP playing the role of BP in the {101} model, with lower invasion time than compartments that require two switchings to get to LP_2_ rather than a single one.

**Figure 5.**
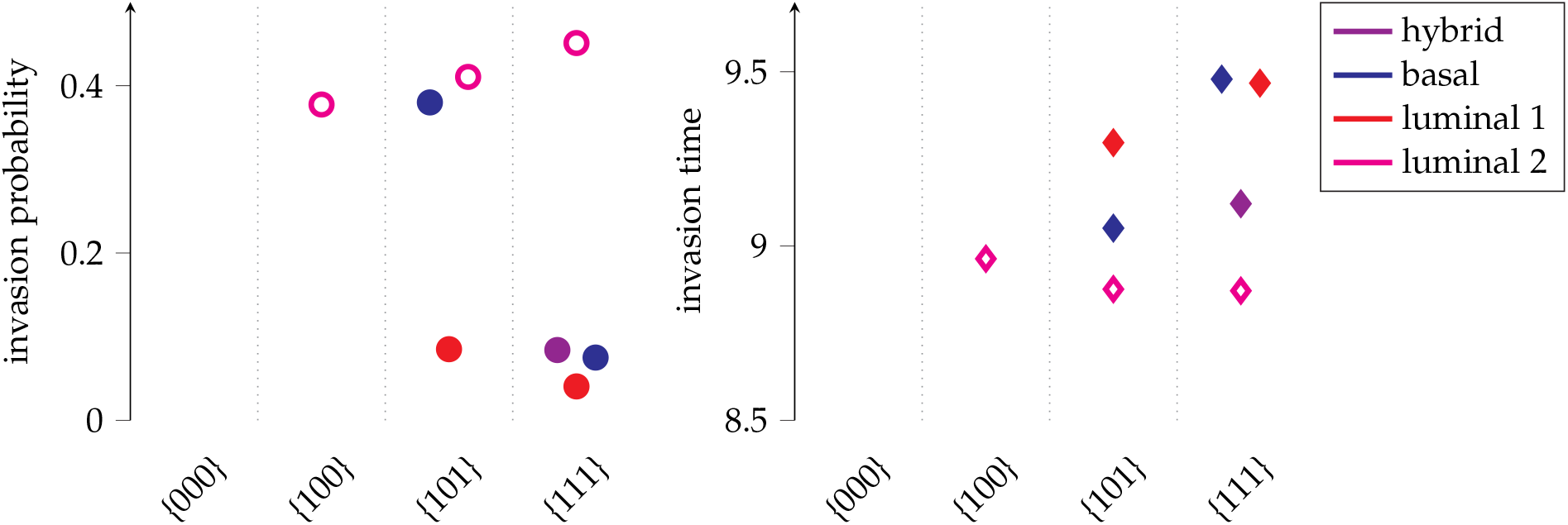
Invasion probability (left panel) and invasion time (right panel; in arbitrary units), averaged over 2000 realisations with real initial populations and fitness *s* = 1, for the four model types. Colours denote which compartment was seeded with a single mutant: basal/BP (blue), luminal 1/LP_1_ (red) or luminal 2/LP_2_ (pink unfilled).

### Invasion, treatment and regrowth population dynamics

Supposing that treatment is imposed once the mutant population has invaded, we can observe population dynamics of the {111} model, as depicted in Figure 6. Here, grey bars denote treatment being applied and solid lines depict mutant population sizes. In addition, pie charts show the distribution of population between the healthy compartments (coloured wedges) and the mutants (black wedges) at three points in time: at the appearance of the mutant, upon invasion, and at the end of the simulation. Two growth dynamics are presented, distinguished by whether or not the mutant population is eradicated by the treatment (left panel) or not (right panel). In both cases, as the mutant population grows, the healthy mature cells are depleted (see dotted lines), since there are fewer healthy progenitor cells (see dashed lines) to maintain them. The right panel then shows an example of a mutant cell acquiring therapeutic resistance, allowing its descendants to invade despite the continued application of treatment. If, instead, all mutants are killed by the first round of treatment as in the left panel, the system returns to homeostatic population sizes. Here, phenotypic switching allows the depleted compartments (where cells of luminal type are severely depleted due to castration [17]) to replenish their populations with the help of other progenitor types via the hybrid phenotype. Finally, note that the mature compartments require more time to reestablish their equilibria population sizes, even with an increased rate of differentiation.

**Figure 6.**
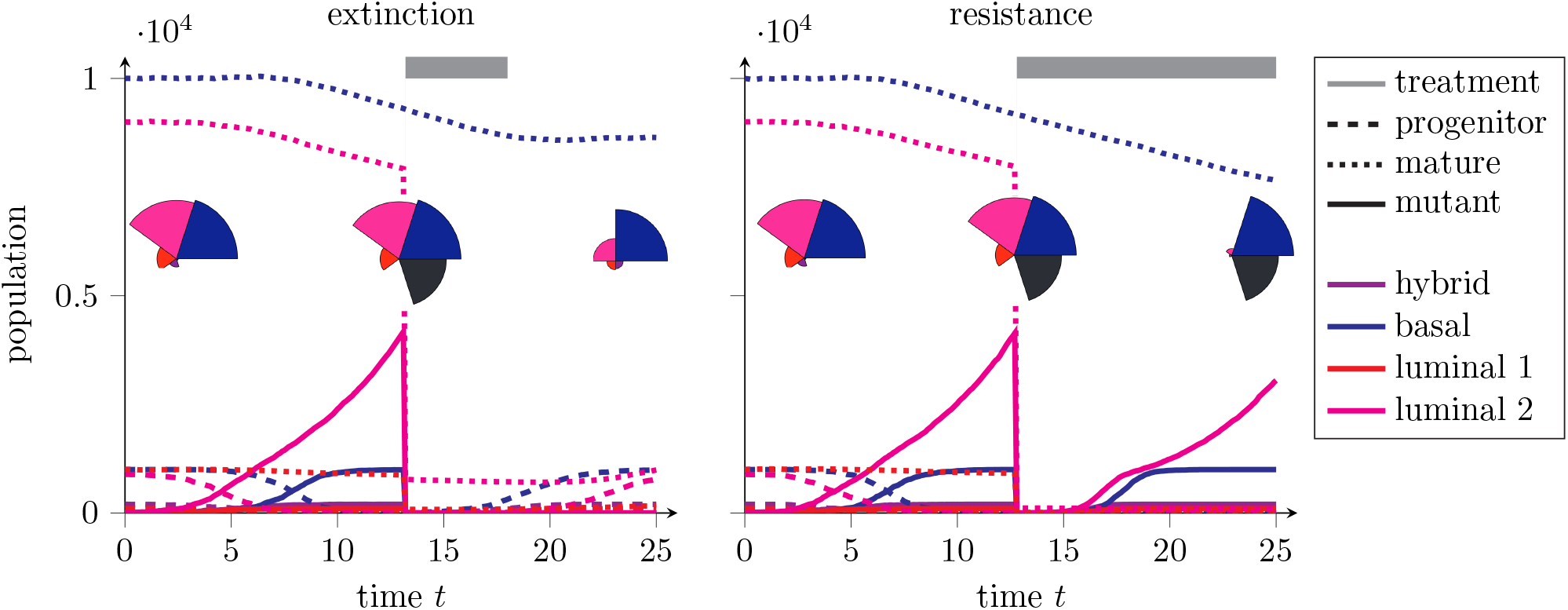
Sample realisations for mutant invasion and treatment (grey bars). All mutant cells may be eradicated with treatment (left), in which case healthy tissue regrows. Or, a resistant subclone survives and invades (right). Population sizes of wild-type progenitor (dashed), mature progenitor (dotted) and mutant (solid) cells in hybrid (purple), basal (blue), luminal 1 (red) and luminal 2 (pink) compartments. Pie charts show the proportion of the population of different phenotypes, with mutants in black.

## Discussion

Multipotent progenitor cells have been observed in and modelled for the skin epidermis, the oesophagus and the liver (see for example [29–32]). In the prostate, however, there is no consensus on the relationship between different cell types and their potencies [15]. The recent evidence of progenitor capabilities for cells in both the basal and luminal compartments suggests a compartmental approach akin to an island model with migration, or a metapopulation model [33]. Using conservation of mass arguments, as done by e.g. Dingli et al. [20], along with making certain assumptions such as equal phenotypic switching rates in and out of each compartment, it is possible to restrict plausible switching, self-proliferation and differentiation parameters. However, these physical restrictions still provide the real organ system with much flexibility when it comes to actual values for these rates. With limited biological data to draw upon for parameterisation, here we explore a stochastic model of the prostate’s compartments, representing the emergence of cancer by a fit mutant.

By allowing the luminal 2 compartment (LP_2_) to have a more flexible population size, assuming some form of less-tight regulation than the other compartments, we witness invasive mutant populations blooming in that compartment (see Figures 3 and 5), aligned with experimental observation [8]. The dependency of the system on the differentiation and switching parameters reveal that the mutant’s relative fitness advantage increased with higher differentiation (see Figure 4), since cancerous mutant cells are assumed to retain their progenitive capabilities. This implies that during periods of more frequent differentiation (such as during wound healing), mutants present in the organ may have an even larger effective fitness advantage than otherwise. Surprisingly, an increase in switching does not directly link to faster invasion times (see Figure 4). Instead, only a minimal presence of phenotypic switching is necessary to induce cancer spread; this aligns with experimental evidence that multipotency in the prostate is linked to tumorigenesis, and tumour heterogeneity is associated to switching rates within the prostate (measured by cell lineage infidelity) [15].

The stochastic switching considered here crucially differs from its deterministic counterpart for the following reason. In a deterministic model with phenotypic switching, mutants arising in different compartments would not have significantly differing invasion times. This is because of the continuous nature of ordinary differential equation models: as soon as a mutant cell appears, it diffuses (fractionally) into other compartments. Thus LP_2_ would, from the time of seeding, have a nonzero mutant population, which would be permitted to grow exponentially immediately. Instead, as shown in the right panels of Figures 3 and 5, the compartment where the mutant was seeded directly impacts the ensuing invasion time. In particular, we can identify two contributions to the invasion time: the time to switch from the seeded compartment to the more loosely regulated compartment (here LP_2_) and the time to grow from a single cell in the less tightly regulated compartment to an invasive population. For any given compartment, the first of these is equal to the sum of the inverse switching rates for each switch required to traverse to LP_2_. This switching rate is equal to the product of the birth rate, the switching probability and the population of mutants in the compartment considered. The second factor can be approximated by the fixation time from a single fit mutant within a compartment undergoing the usual Moran process, which has been calculated under certain conditions (see for example [34–36]).

Mutant extinction is also stochastic, where extinctions occur early and mostly independently from the compartment of origin (see Figures S4-S6). The exception to this is the hybrid compartment (HP), where mutant cells seeded there are more likely to go extinct. This is because other cell types undergo more switching events into HP, possibly replacing the mutant, than the other progenitor types, which only admit phenotypic switches from HP alone.

Though the multipotent hybrid progenitor cell type is imprecisely defined in terms of biology [18], it seems to play a more significant role under non-homeostatic conditions [8, 15, 16]. In particular, under injury (such as luminal ablation due to androgen deprivation), the hybrid cells play a role in regenerating the tissue, before decreasing in prevalence as the tissue returns to normal [15]. The first of these observations is present in our model results, where the phenotypic switching allows for both the spread of the mutant (Figure 5) but also for the replenishing of underpopulated compartments (Figure 6). A shortcoming of our model is its inability to incorporate flexible compartment sizes: HP is present throughout, rather than appearing when needed and fading when not. Inclusion of a dynamic network topology would allow for this plasticity, but it is beyond the scope of this work.

While we mostly focus on invasion probabilities and times, several conclusions can be drawn from the population dynamics. Figure 6 demonstrates that upon cancer emergence, the basal population decreases, even though the mutant population only explodes in the luminal compartment. This is supported by some experimental results, which suggest that basal cell depletion is a marker for prostate cancer [8]. Our results provide a mechanistic explanation for this phenomenon: with fewer healthy basal progenitors (since a subset has become cancerous), the mature population that can be maintained diminishes. This depletion occurs no matter which compartment was seeded, since even when the cancer originates in a non-basal compartment, the tumour will spread to the basal compartment.

Finally, as evidenced by Figure 6, inherent to compartmental models with phenotypic switching is treat-ment failure whenever the treatment does not target all cell types and immediately eradicate all cancer cells directly. To investigate this, we modelled androgen deprivation therapy, a treatment option that primarily targets luminal cells, which are dependent on hormones for growth and survival [5]. Because at the time of treatment there are cancer cells in the basal compartment, these go unharmed by ADT and thus allow for the tumour to regrow once therapy ends by transitioning back into a luminal state. This is in direct contrast to the perspective that basal cells obstruct oncogenic offenses from impacting the luminal compartment—by acting as upstream progenitors of luminal cells, as well as by physically forming a blood-prostate barrier, allowing them to regulate functions of the luminal cells—as has been suggested [7]. In addition, selection for treatment-resistant cells takes place, which is one of the primary causes of hormonal treatment failure and tumour progression [37]. This sheltering effect reinforces that for elimination of prostate cancers, a combination of multiple forms of therapy may be necessary, such as radiotherapy, chemotherapy, or immunotherapy in addition to ADT.

## Acknowledgements

This research was supported by the European Union’s Horizon 2020 research and innovation programme under the Marie Skłodowska-Curie EvoGamesPlus grant number 955708.

