## Extended Figures for "Stochastic modelling of prostate progenitor architecture"

### Supplementary Information

If smaller compartment populations are used for the simulations, then similar patterns are observed. For example, in Figure S1, initial population sizes for HP, BP, $LP_1$ and $LP_2$ are equal to 10, 15, 20 and 50, respectively. (Again, the progenitor population sizes account for 10% of the mature population size for each compartment.) Here, it is clear that mutants seeded in $LP_2$ have the highest invasion probability and the quickest invasion time. We once more have that increasing the fitness allows one to distinguish the number of switchings required to get from the seeded compartment to $LP_2$.

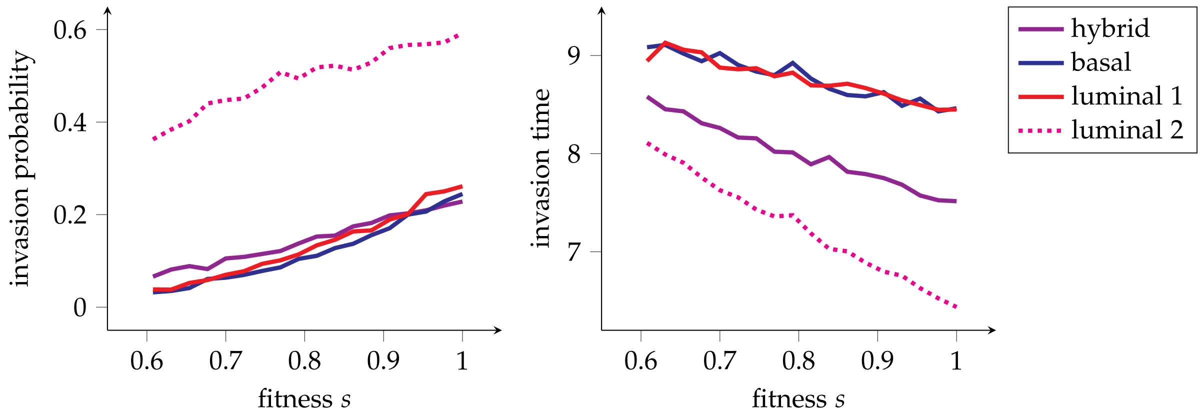


**Figure S1.** Invasion probability (left panel) and invasion time (right panel; in arbitrary units) versus fitness, averaged over 3000 realisations, where compartment populations are small (see text for details). Colours denote which compartment was seeded with a single mutant: hybrid/HP (purple), basal/BP (blue), luminal 1/$LP_1$ (red) or luminal 2/$LP_2$ (pink dotted).

Figure S2 depicts the corresponding plots to Figure 3 for the other models ({100} and {101}), where model {000} is not present because no invasion takes place.

Figure S3 shows the same heat maps as Figure 4 but for mutants seeded into $LP_1$, exhibiting the same behaviour as for BP (middle row of Figure 4).

Figure S4 demonstrates the relationship between extinction probabilities and times with fitness. In particular, with greater fitness, the likelihood and speed of extinction decrease, unsurprisingly. This behaviour in general is shared by all compartments, in all models.

There are two contributions to the higher extinction probability and shorter extinction time experienced by mutants seeded in HP of the {111} model (rightmost column of Figure S4): first, a relatively small population, so upon a switching event into the compartment, a randomly selected cell to be replaced is more likely to be the mutant. That is, for a neighbouring compartment that is five times more populous such as BP, switching events into HP are five times more frequent, so the mutant population in the smaller compartment is more likely to be replaced by the immigrating cells. Second is the number of switches itself, captured by the number of compartments to which a compartment is connected via phenotypic switching. In the case of HP, this is three (BP, $LP_1$ and $LP_2$), whereas the other three only have one connection (to HP). This surplus of switching events exacerbates the first effect. In the {111} model, $LP_1$ only experiences the first of these effects, so has higher extinction probability than BP and $LP_2$, as seen in the upper-right plot of Figure S4. In the {101} model, $LP_1$ also faces the first of these effects, but now its neighbouring compartment is much larger (BP is a factor of five larger than HP), so the effect is accentuated, leading to an elevated extinction probability. While BP experiences the second effect (as it is connected to both $LP_1$ and $LP_2$), its impact is not significant enough to be discerned in Figure S4. Note finally that the extinction times are much lower than the invasion times seen in previous figures; this demonstrates that mutants only die out early, by stochastic extinction, whereas growing to a detectable size requires more time to elapse.

Next, Figure S5 presents heat maps for varying differentiation and switching probabilities of the extinction probability and time for mutants seeded in different compartments. Like the invasion probability and time, different patterns emerge depending on the compartment of origin: HP undergoes more and faster extinction for higher switching probabilities (first row); more differentiation corresponds to higher relative mutant fitness and therefore lower extinction probability (second row); $LP_1$, being smaller than the others (and thus experiencing more significant small population stochastic effects), faces both HP- and BP-type impacts to its extinction probability (third row); and $LP_2$ behaves as BP, with no impact of switching (fourth row).

Finally, Figure S6 depicts the extinction probabilities and times for different models; here, identical effects to those shown in Figure S4 are present.

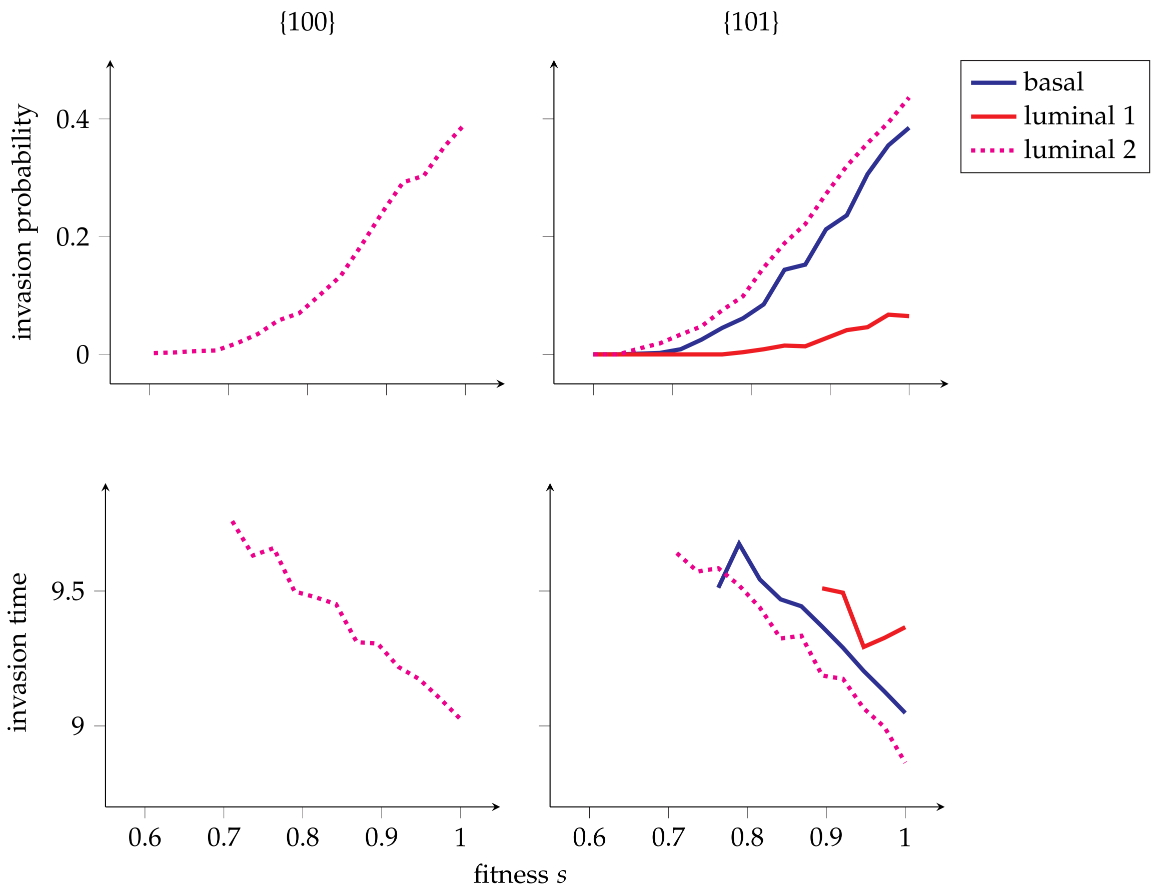


**Figure S2.** Invasion probabilities (top) and invasion times (bottom; in arbitrary units) versus fitness for the {100} (left) and {101} (right) models, averaged over 100 realisations. (Note that no invasion took place in the {000} model.) Colours denote which compartment was seeded with a single mutant: basal/BP (blue), luminal 1/$LP_1$ (red) or luminal 2/$LP_2$ (pink dotted).

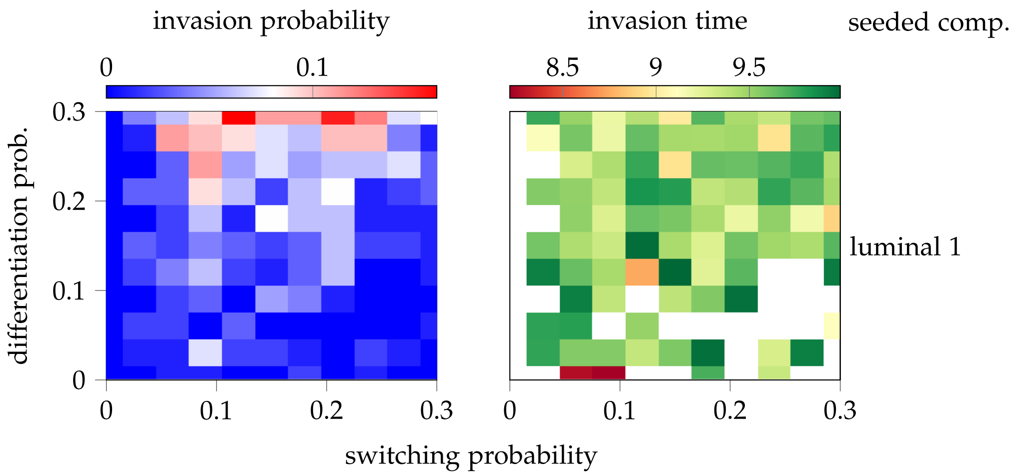


**Figure S3.** Invasion probabilities (left) and invasion times (right; in arbitrary units) for different switching and differentiation probabilities when the mutant is seeded in the luminal 1/$LP_1$ compartment. Each cell is averaged over 100 realisations; blank cells in the right column indicate that zero invasions took place.

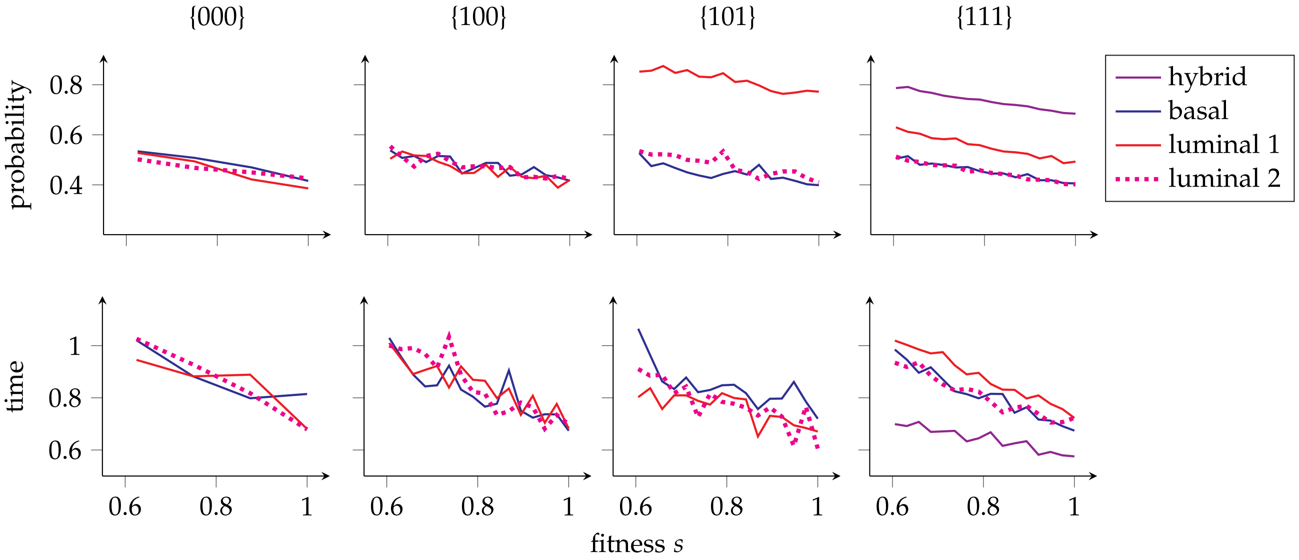


**Figure S4.** Extinction probabilities (top) and extinction times (bottom; in arbitrary units) versus fitness for all four models, averaged over 1000 realisations (except for {111}, which was averaged over 5500 realisations). Colours denote which compartment was seeded with a single mutant: basal/BP (blue), luminal 1/$LP_1$ (red) or luminal 2/$LP_2$ (pink dotted).

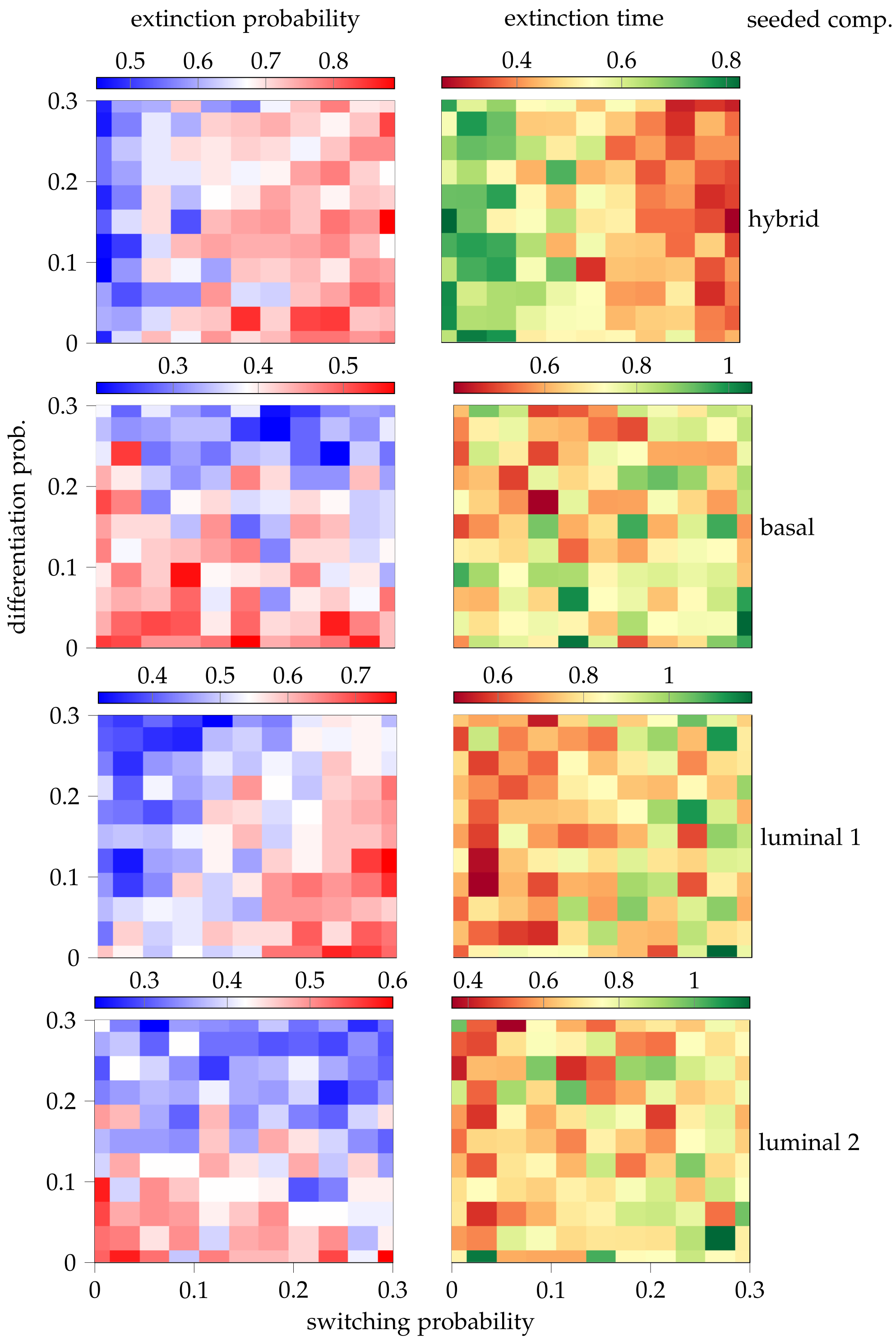


**Figure S5.** Extinction probabilities (left) and extinction times (right; in arbitrary units) for different switching and differentiation probabilities when the mutant is seeded in each compartment. Each cell is averaged over 100 realisations.

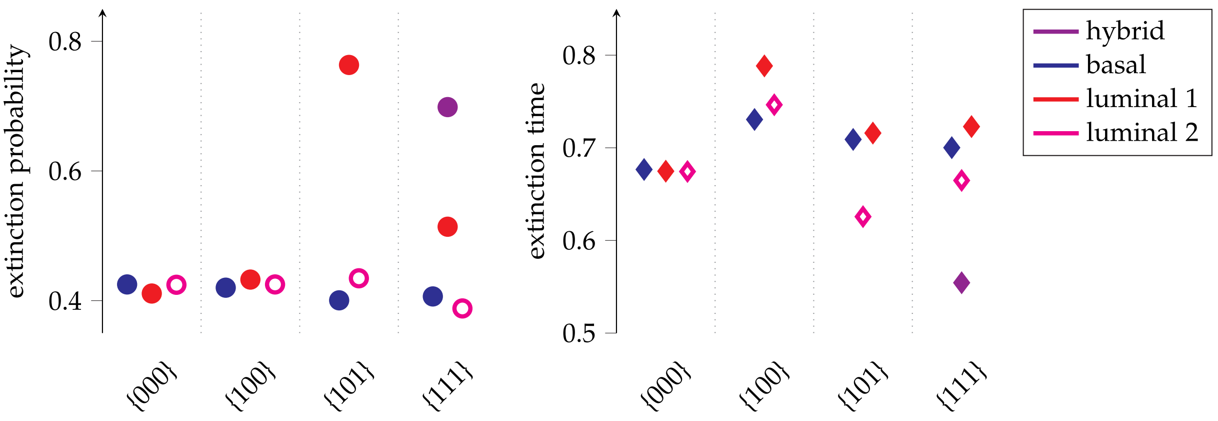


**Figure S6.** Extinction probability (left panel) and extinction time (right panel; in arbitrary units), averaged over 2000 realisations with fitness $s = 1$, for the four model types. Colours denote which compartment was seeded with a single mutant: basal/BP (blue), luminal 1/$LP_1$ (red) or luminal 2/$LP_2$ (pink unfilled).
